# Molecular Inverse Comorbidity between Alzheimer’s disease and Lung Cancer: new insights from Matrix Factorization

**DOI:** 10.1101/643890

**Authors:** Alessandro Greco, Jon Sanchez Valle, Vera Pancaldi, Anaïs Baudot, Emmanuel Barillot, Michele Caselle, Alfonso Valencia, Andrei Zinovyev, Laura Cantini

## Abstract

Matrix Factorization (MF) is an established paradigm for large-scale biological data analysis with tremendous potential in computational biology.

We here challenge MF in depicting the molecular bases of epidemiologically described Disease-Disease (DD) relationships. As use case, we focus on the inverse comorbidity association between Alzheimer’s disease (AD) and lung cancer (LC), described as a lower than expected probability of developing LC in AD patients. To the day, the molecular mechanisms underlying DD relationships remain poorly explained and their better characterization might offer unprecedented clinical opportunities.

To this goal, we extend our previously designed MF-based framework for the molecular characterization of DD relationships. Considering AD-LC inverse comorbidity as a case study, we highlight multiple molecular mechanisms, among which the previously identified immune system and mitochondrial metabolism. We then discriminate mechanisms specific to LC from those shared with other cancers through a pancancer analysis. Additionally, new candidate molecular players, such as Estrogen Receptor (ER), CDH1 and HDAC, are pinpointed as factors that might underlie the inverse relationship, opening the way to new investigations. Finally, some lung cancer subtype-specific factors are also detected, suggesting the existence of heterogeneity across patients also in the context of inverse comorbidity.

## 1. Introduction

Large-scale genomics projects, including for instance The Cancer Genome Atlas (TCGA, https://www.cancer.gov/tcga), are currently providing an overwhelming amount of omics data. The available data offer the opportunity to better understand biological systems and cancer in particular, but their high dimensionality poses considerable challenges typical of “Big Data” [1].

A powerful approach to this problem is represented by Matrix Factorization (MF), a class of unsupervised methods that reduces high-dimensional data into low dimensional subspaces, while preserving as much information as possible [2–4]. Given a data matrix χ, MF learns two sets of low-dimensional representations: “metagenes”, encoding molecular relationships, and “metasamples”, encoding sample-level relationships. Up to now, MF has been successfully used in a broad spectrum of applications: unsupervised clustering, especially in the context of cancer subtyping [5,6], molecular pattern discovery [7,8], mutational signatures definition [9,10] and tumour sample immune infiltration quantification [11]. Such results have been obtained by mining with MF single large-scale datasets, such as transcriptome or methylome. Recently, we designed a metric to infer univocal correspondences between the metagenes obtained by an MF algorithm on multiple independent datasets profiled from the same biological condition (e.g. same cancer tissue), and used this metric to design a methodological framework that revealed relevant pathways characteristic of colorectal cancer [12].

We are here interested in investigating the molecular bases of previously documented Disease-Disease (DD) relationships. Indeed, several computational studies have inferred DD relationships, starting from the “Human Disease Network” where diseases were connected when sharing disease genes [13], to the “multiplex network of human diseases” composed by genotype- and phenotype-based layers that propose new disease-associations [14]. More importantly, DD relationships have also been systematically identified by epidemiological studies, working at the level of populations and looking for the co-occurrence of different diseases in the same patients by using medical claims [15], medical records [16] and insurance claims [17]. The higher than expected risk of developing pancreatic cancer in patients suffering for type II diabetes [18] and of developing lung cancer in asthma patients [19] are among the most renown examples of cancer-related comorbidities. Interestingly, it has also been described that patients suffering from certain diseases have a lower than expected risk of developing cancer, known as inverse comorbidity [20–22]. An example of these protective effects of one disease on the other is represented by the documented inverse comorbidity between Alzheimer’s Disease (AD) and Lung Cancer (LC) [22–24]. Molecular and non-molecular factors (e.g. the environment, lifestyle or drug treatments) can be responsible for such DD relationships. The molecular mechanisms underlying these DD relationships are poorly understood and investigating them offers unprecedented opportunities to better understand the etiology and pathogenesis of diseases, with the hope of identifying opportunities for repositioning of pre-existing treatments.

Recently, transcriptomic meta-analyses revealed sets of significantly up and down regulated genes that are shared across diseases displaying different patterns of direct and inverse comorbidities [25,26]. However, differential expression analysis only focuses on the predominant signals present in the data, failing to capture alternative signals and local behaviors [3]. These limitations are overcome by MF that learns metagenes, i.e. ranking of genes, without focusing on single sets of predominant genes. Moreover, contrarily to differential expression analysis, MF jointly provides metagenes and metasamples, i.e. also grouping samples together with their biological characterization. We hereby propose to use an MF approach to study the molecular bases of DD relationships. This, however, requires innovative adaptations. We thus propose to extend our previously defined MF framework for the particular study of DD relationships [12]. Moreover, given the existence of positive and negative DD connections, we also adapt the framework to distinguish molecular relationships concordantly and discordantly altered in datasets coming from different diseases.

Considering the inverse comorbidity between Alzheimer’s disease (AD) and lung cancer (LC) as a case study [22–24], we applied our MF framework to 17 transcriptomic datasets, including both LC and AD samples (total of 1367 samples), and we highlighted multiple molecular mechanisms possibly underlying the inverse comorbidity pattern. Through a pancancer analysis we categorized the processes here suggested to be involved in the AD-LC inverse comorbidity based on their presence in other cancers. The previously identified role of the immune system and mitochondrial metabolism in AD-LC inverse comorbidity is confirmed by our analysis. Additionally, new candidate molecular players, such as Estrogen Receptor (ER), CDH1 and Histone Deacetylase (HDAC), are identified as potentially involved in the inverse comorbidity considered. Finally, some lung cancer subtype-specific alterations are also detected suggesting the existence of heterogeneity across patients also in the context of inverse comorbidity.

## 2. Results

### 2.1. A new MF framework to study disease-disease relationships

We previously defined the Reciprocal Best Hit (RBH) metric to infer univocal correspondences between the MF metagenes obtained on independent datasets measured from the same biological condition (e.g. same cancer tissue) [12]. Based on this metric we designed an RBH-based framework, structured in three sequential steps: (1) each transcriptomic dataset is independently decomposed in metagenes and metasamples with MF; (2) using the RBH metric, relationships between metagenes are inferred and a RBH network is constructed; (3) communities are detected in the RBH network. These communities of genes are then analyzed for functional relatedness and provide a biological interpretation of the principal factors that shape the transcriptomes. Here, we adapted the framework to the study of the molecular mechanisms underlying DD relationships, in order to infer univocal positive/negative correspondences between MF metagenes independently obtained on datasets measured from different diseases. Briefly, the main methodological novelties are: (i) the investigation of a methodology for the orientation of the metagenes (i.e. assign a sign to the metagenes, in order to express either direct or inverse similarity between them); (ii) a novel definition of Reciprocal Best Hit (RBH) network taking into account the orientation of the metagenes and (iii) the restriction of the community detection phase to the subnetwork of interest (e.g. subnetwork of negative links connecting metagenes of LC and AD in our case). The structure of the framework together with its novelties is summarized in Figure 1.

**Figure 1.**
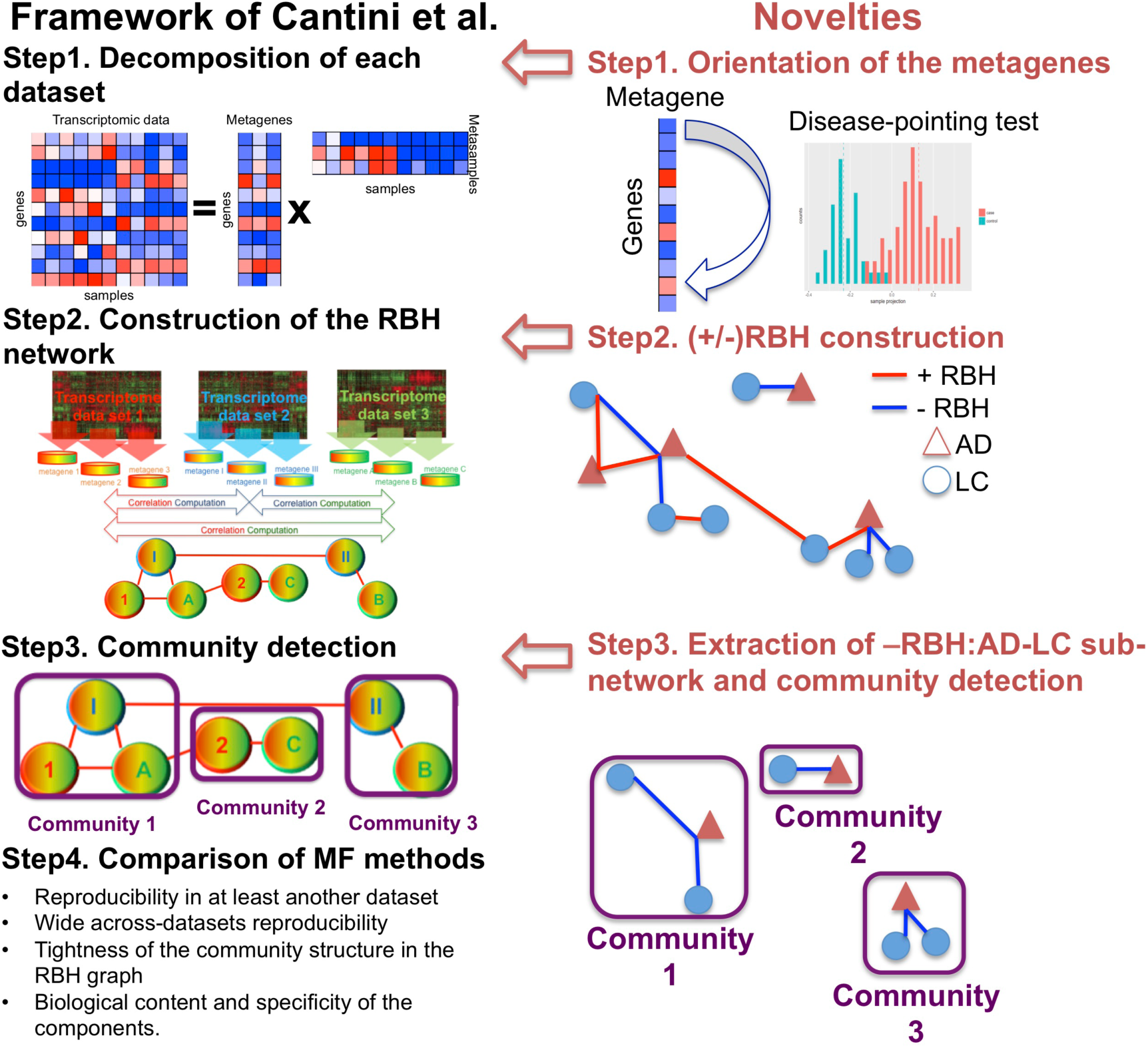
Schematic view of the framework and the novelties introduced with respect to [12].

#### 2.1.1. Setp1: data decomposition and orientation of the components

Each transcriptomic datasets is separately decomposed using MF. The framework here proposed can be combined with the MF algorithm of interest. In this work, we chose stabilized Independent Component Analysis (sICA) [12,27,28], a stabilized version of ICA [28–30]. SICA was indeed previously shown to outperform alternative MFs in the extraction of relevant biological knowledge from collections of transcriptomic datasets derived from the same biological condition (e.g. the same cancer type) [12]. Moreover, the ability of sICA to separate the various overlapping biological factors present in transcriptomic data, such as those linked to the tumor cells, the tumor microenvironment and non biological factors, linked to sample processing or data generation, makes this approach particularly promising for extracting relevant molecular factors from the numerous confounding factors involved in DD relationships.

By applying sICA to a transcriptomic matrix *χ (n x m)*, with *n* genes in the rows and *m* samples in the columns, we reduce it to the product of an unknown mixing matrix *A (n x k)*, whose columns are here denoted as “metagenes” and an unknown matrix of source signals *S (k x m)*, whose rows are here denoted as “metasamples”. The metagene/metasample associated to the component *i* will thus provide the contribution of each gene/sample present in the matrix *χ* to component *i*. Metasamples and metagenes are learned based upon the assumption that the number *k* of components occurring in the input matrix *χ* is smaller than either its rows or columns. We here selected the number *k* of components equal to 100 for those datasets having more than 100 samples and equal to half of the samples for smaller datasets. These chosen values are higher than the estimation of the optimal transcriptomic dimension, due to the fact that overdecomposition in sICA was proven not detrimental for the interpretability of the resulting components [28].

To determine the orientation of the sICA metagenes two alternative approaches are considered: “Long tail-pointing” and “disease-pointing”. The long tail-pointing approach, previously used for other sICA applications [28], orients the metagenes such that the longest tail of their distribution corresponds to their positive side. Indeed the sICA factors are identified by maximizing non-gaussianity of the data point projection distributions. As a consequence, the longest-tails of such distributions are those containing most of the biological information. We here introduce the “disease-pointing” approach, which exploits the availability of cases and control samples to orient the components. More specifically, the differential association of each metasample to case vs. control is tested based on a Wilcoxon test and the couple metagene-metasample is oriented so that the cases in the metasample are on the positive side.

#### 2.1.2. Step 2: Construction of the signed Reciprocal Best Hits (sRBHs)

At Step 2, the Reciprocal Best Hit (RBH) network is constructed. A positive/negative RBH is defined as follows: given two sets of metagenes {*M*_*1*_…*M*_*k*_} and {*N*_*1*_…*N*_*k*_} obtained from the transcriptomic datasets *T*^*M*^ and *T*^*N*^, respectively, we define *M*_*i*_ and *N*_*j*_ a positive Reciprocal Best Hit (+RBH) iff

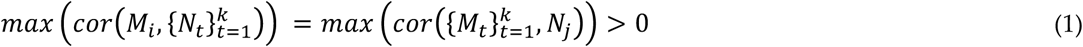

and *M*_*i*_ and *N*_*j*_ a negative Reciprocal Best Hit (-RBH) iff

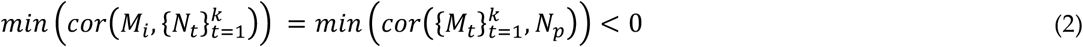

Each metagene will thus find a maximum of two associated metagenes in another independent transcriptomic dataset, corresponding to +RBH (1) and -RBH (2). Repeating the same procedure for the metagenes of all the available transcriptomes we obtain a network whose nodes are the metagenes computed in all the transcriptomic datasets and whose links correspond to their +RBH and -RBH computed as in (1,2).

#### 2.1.3. Step3: subnetwork isolation and community detection

In step 3, given our interest for the processes that are differentially altered between two diseases, such as AD and LC, we delineate the relevant subnetwork of RBHs. For example, if we want to study the inverse comorbidity between AD and LC, we restrict the analysis to the negative RBHs (-RBHs) connecting metagenes of AD with metagenes of LC. Once selected the subnetwork of interest, we detect communities with the MCL algorithm [31,32]. Such communities correspond to highly reproduced biological components involved in DD relationship. Moreover, having previously isolated the subnetwork of interest (such as negative RBHs between AD and LC) we are sure to only identify communities that are altered in the same direction of the comorbidity under analysis (oppositely regulated in case of inverse comorbidity and concordantly regulated in case of positive comorbidities). The obtained communities are then biologically annotated and interpreted as described in Methods.

### 2.2. Investigation of the orientation methodology for the sICA components

Among the various modifications apported to the framework, of particular importance is the choice of the procedure for the orientation of the metagenes. As described previously, two alternative approaches were considered: “Long tail-pointing” and “disease-pointing”. We tested how such choice impacts the following steps of the framework and, in particular, the structure of the obtained RBH network. To do this, we selected a specific DD relationship, i.e., the inverse comorbidity between Alzheimer’s disease (AD) and lung cancer (LC) as a case study [22–24].

17 transcriptomic datasets, spanning AD and LC patients and containing case and control samples, were employed (see Methods for further details). Following our framework (Figure 1), each dataset was decomposed separately through sICA (see Supp Table 1 for the number of components) and the orientation of the components was established both with the long-tail-pointing and the disease-pointing approaches. The resulting metagenes were then compared according to multiple criteria (see Figure 2).

**Figure 2.**
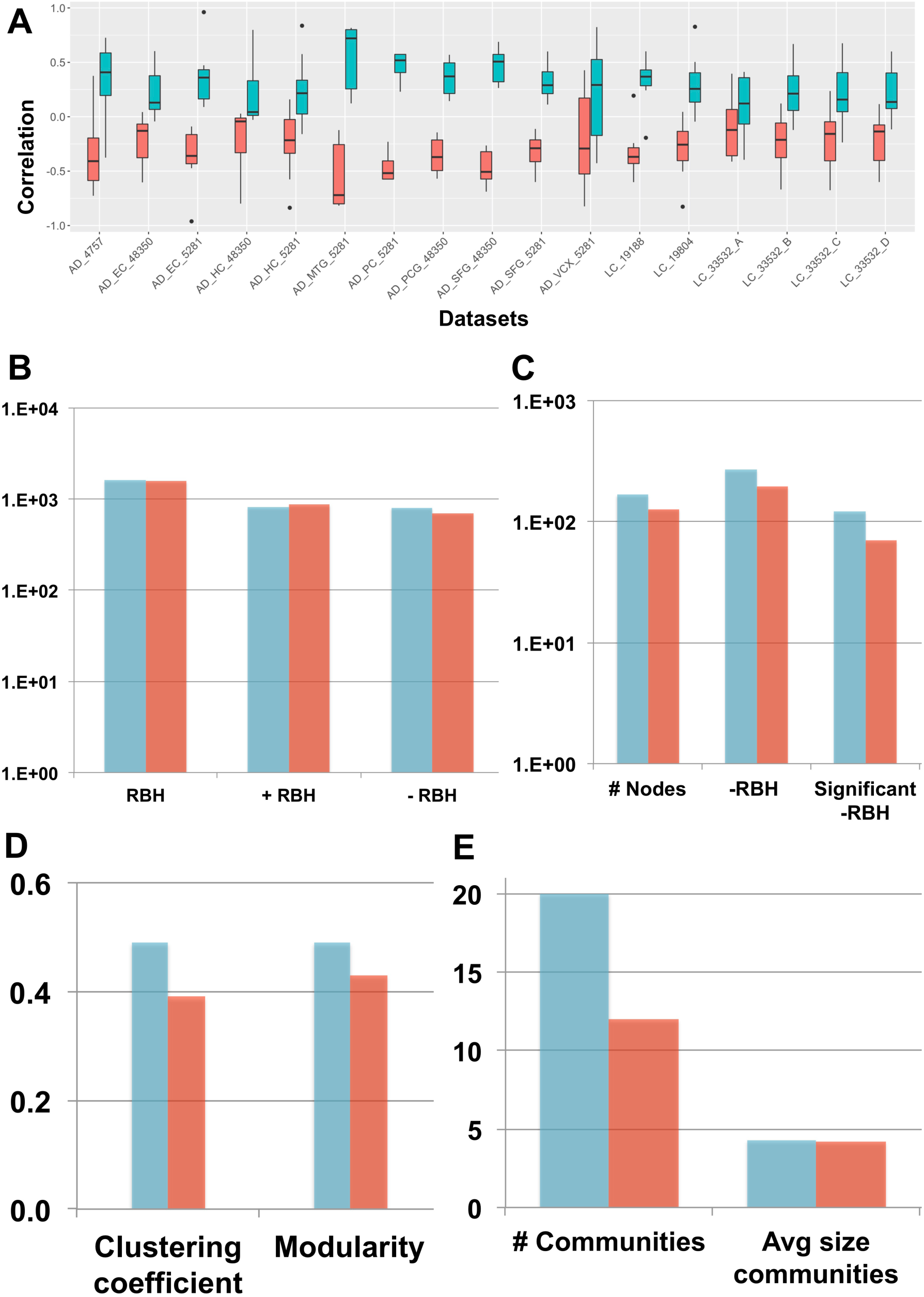
“Long-tail-pointing” (red) vs. “disease-pointing” (blue) orientation of the sICA factors. (A) The two methods of factors orientation are compared based on the correlation of the obtained metagenes with the case vs. control genes’ fold change. (B) The two methods are compared based on the number of links present in their RBH network. Total RBHs (RBH), positive RBHs (+RBH), negative RBHs (-RBH). (C,D,E) The two methods are compared based on the structure of their -RBH AD/LC subnetwork, relevant for the study of inverse comorbidity. In (C) the number of nodes and links in the subnetwork are compared. Number of nodes (# Nodes), negative RBHs connecting an AD component with a LC component (-RBH) and negative RBHs connecting an AD component with a LC component that are associated to nodes with significant differential behaviour (Wilcoxon p-value < 0.05) between case and control (significant -RBH). In (D) The clustering coefficient and modularity of the subnetwork are considered. In (E) The number of communities and their average size in the subnetwork is taken into account.

First, the correlation between the obtained metagenes and the case vs. control fold-change of expression was considered. In fact, to associate a metagene to a specific biological function or pathway, we need to perform enrichment tests using databases of functional annotations (e.g. Reactome, GO). Generally this interpretation step is just aimed at associating a function to each metagene, without taking into account the sign of activity of the identified pathways/processes. However, when dealing with comorbidities it is important to not only associate a function to each metagene, but also to infer the sign of activity of such pathways/functions. This task can be easily achieved once the metagenes are positively correlated with the gene fold-change. As shown in Figure 2A, the disease-pointing orientation produces metagenes that are significantly more correlated with the genes fold-change than the long tail-pointing one (significance tested with Wilcoxon test, resulting P-values available in Supp Table 1).

We have then applied the Step 2 of the framework and independently constructed an RBH network for “long-tail-pointing” and “disease-pointing” oriented metagenes. In both cases, the nodes of the network correspond to the metagenes independently identified in the 17 datasets (369 total nodes) and their links are +/-RBHs, defined as in equations (1,2). Changes in the orientation of the metagenes alter the sign of the correlations giving rise to different RBH networks. We have thus compared the “long-tail-pointing” vs. “disease-pointing” RBH networks based on their number of links (Figure 2B). The “disease-pointing” method returns 1616 RBHs vs. the 1574 returned by the “long-tail-pointing” method. Such result is due to the higher number of -RBHs identified with the “disease-pointing” orientation (802 vs. 705).

In Step 3 we focused on the subnetwork composed of -RBHs and linking AD components with LC ones and vice-versa, which in the following we call “-RBH AD/LC subnetwork”. These are in fact the metagenes and RBHs of interest for the study of AD-LC inverse comorbidity. We studied the topology of this subnetwork starting from its number of nodes and links (Figure 2C). The -RBH AD/LC subnetwork based on the “disease-pointing” orientation includes a higher number of metagenes (167 vs. 127 of “long-tail-pointing”) and a higher number of links (268 vs. 194 of “long-tail-pointing”). Moreover, among the RBHs present in the subnetwork, those of the “disease-pointing” tend to be more frequently connecting factors that are significantly differential between case and control (112 vs. 70 of “long-tail-pointing”). Communities were then detected in the “long-tail-pointing” and “disease-pointing” x-RBH AD/LC subnetworks. As shown in Figure 2D, the “disease-pointing” -RBH AD/LC subnetwork has a higher modularity (0.49 vs. 0.43) and higher clustering coefficient (0.49 vs. 0.39). Moreover 20 communities of size higher or equal to 4 are detected in the “disease-pointing” -RBH AD/LC subnetwork vs. the 12 of the alternative approach and the average size of the “disease-pointing” communities is 4.3 vs. the 4.2 of the alternative approach (Figure 2E).

Overall our analysis indicates that the “disease-pointing” orientation tends to identify a higher number of candidate molecular processes/pathways involved in AD-LC inverse comorbidity. For all these reasons, “disease-pointing” is the orientation approach that we selected for the following analysis.

### 2.3. New biological insights on the inverse comorbidity between AD and LC

We hypothesize that the communities of the -RBH AD/LC subnetwork, obtained with the “disease-pointing” orientation, could be related to the AD-LC inverse comorbidity. We thus annotated the communities of the -RBH AD/LC subnetwork by using MsigDB signatures [33], Microenvironment Cell Populations-counter (MCP-counter) signatures [34], predefined lung cancer subtypes [35] and the metagenes computed in [27], here referred to as CIT, as described in Methods. The obtained -RBH AD/LC subnetwork with the main biological information is illustrated in Figure 3 and Supp Table 2.

**Figure 3.**
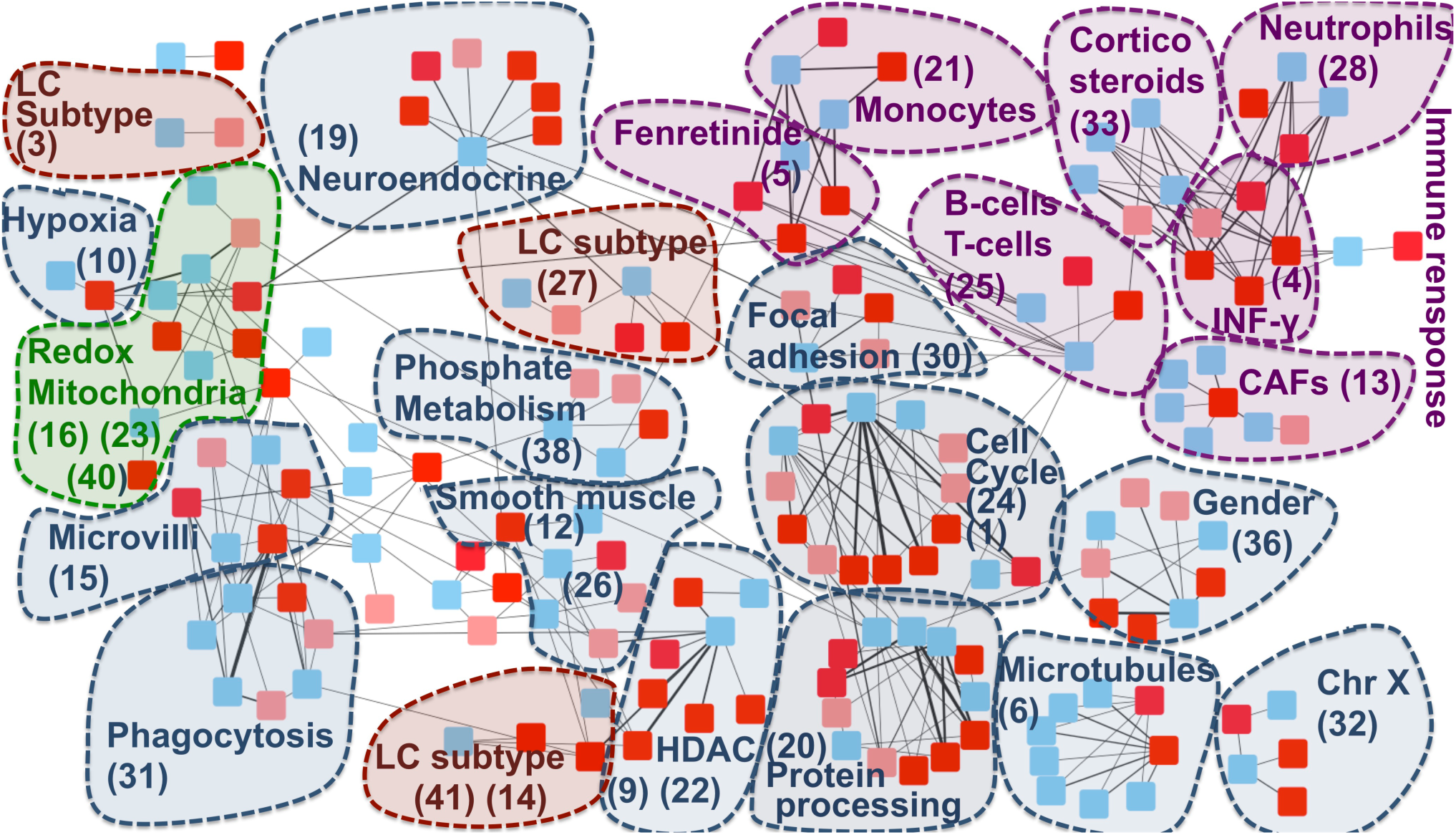
RBH AD/LC subnetwork with biological annotations. Each node in the network corresponds to a metagene, the list of metagenes associated to each community ID is reported in Supp Table 3. Colours are linked to the diseases: red for AD and blue for LC. In AD datasets obtained from the same region of the brain are denoted with different shades of red (normal and light red). The nodes are organized into communities. Each community is denoted with a number corresponding to its ID and the main biological annotation associated to them (see Supp Table 2 for an extensive report).

The majority of the communities present in the network are associated to the immune system and mitochondrial functioning, confirming the results of previous transcriptomics meta-analyses on the inverse comorbidity between AD and LC [25,26]. Interestingly, these processes are here deeply partitioned into multiple communities, suggesting that we can detect more detailed aspects of their involvement. Fibroblasts, Neutrophils, Monocytes, B and T cells are the immune cells showing an inverse activity in LC and AD according to our analysis. Moreover, communities involved in the regulation of two immune-system related drugs (fenretinide and corticosteroids) are identified. Interestingly, corticosteroids are associated with less Alzheimer neuropathology [36], while their use in LC patients is associated with lower overall survival [37]. At the same time, fenretinide has been shown to inhibit growth in lung cancer cell lines [38] and it has been proposed as a potential adjuvant for late onset Alzheimer’s disease [39]. Moreover, the fenretinide community is tightly linked with the monocytes one, in agreement with its mechanisms of action involving the regulation of the secretion of pro-inflammatory cytokines in human monocytes [40].

The communities associated to mitochondria span different processes related to their activity: oxidation-reduction process (communities 16, 23, 40), hypoxia (community 10) and phosphate metabolic process (community 38). Enrichment in hypoxia could correspond to a confounding factor linked to the state of the profiled tissues (post-mortem for AD and fresh tissue biopsy for LC). However, patients suffering from systemic or prenatal hypoxia have a higher risk of developing Alzheimer’s disease [41,42] and targeting hypoxia seems to Improve lung cancer outcome [43], indicating that such hypoxia-related community could also contain non-trivial information.

Additionally to mitochondria and immune system, community 36 has been associated to gender, in line with the higher risk of females to develop Alzheimer’s disease, in opposition to lung cancer, which is more frequent in men [44,45]. Histone Deacetylase (HDAC), associated to community 22, confirms the known involvement of HDAC1 in both cancer and Alzheimer’s disease [46,47]. Community 30 is enriched in focal adhesion. The inhibition of focal adhesion kinase, which is overexpressed in several cancers, decreases cell viability [48], while, in the case of Alzheimer’s disease, amyloid-ß induces the inactivation of focal adhesion kinase [49]. Cell cycle and CDH1 targets are associated to community 24. Interestingly, growing evidence suggests that dysregulation of APC/C-Cdh1 is involved in neurodegenerative diseases, potentially as a consequence of amyloid-ß driven proteasome-dependent degradation of CDH1 [50]. On the other hand, significantly higher methylation level of CDH1, inducing its inactivation, plays an important role in lung cancer [51]. Community 20 is associated to protein processing and chaperone-mediated protein folding. Protein misfolding is a known marker of AD [52,53]. At the same time, cell division, migration, and invasion rely on microtubules and actin filament components and thus chaperone-mediated protein folding activity is tightly linked to cancer [54]. Similar arguments support the involvement of microtubules (community 6) to AD-LC inverse comorbidity. Moreover, response to Estrogene Receptor (ER) (“ESR1 targets”) has been found enriched in 20 communities, even if without clear association to a specific one. Interestingly, an inverse association has been shown between the use of estrogen and early onset of Alzheimer’s disease, suggesting that it might be beneficial for the disease [55], potentially due to its inhibitory activity on neuroinflammation [56]. On the other hand, the use of hormonal replacement therapy significantly increases LC mortality, supporting a role of estrogen in lung cancer [57]. These inverse effects of regulation of focal adhesion, CDH1 and estrogen receptor in cancer and AD are consistent with a possible association of these pathways to the inverse comorbidity patterns observed.

We then explored if also lung cancer subtype-specific molecular mechanisms could be involved into the AD-LC inverse comorbidity [35]. The main biologically-annotated network communities resulted to be general of LC with no association to a specific subtype. At the same time, three communities in our network (27, 14 and 3) mapped to the three predefined LC subtypes (proximal proliferative, proximal inflammatory and terminal respiratory unit, respectively). Therefore, some lung cancer subtype-specific regulatory programs seem to also be involved, suggesting the existence of an across-patients heterogeneity, even if such phenomenon is not a predominant one.

Finally, AD has been shown to have comorbidity relationships at the epidemiological level not only with LC, but also with other cancer types, with most of them being inverse comorbidities [13]. We thus tested if some of the candidate biological processes that we here identified to be possibly involved into AD-LC inverse comorbidity could be generalized to the comorbidity relationship between AD and other cancers. With this aim, we considered metagenes previously computed on TCGA transcriptomes for 32 different cancer types (pancancer metagenes) [28] and inferred their RBHs with the metagenes of the -RBH AD/LC subnetwork. The presence of pancancer metagenes in the communities of the -RBH AD/LC subnetwork has then been tested. If a community in the -RBH AD/LC subnetwork is found to be correlated with some pancancer metagenes we can infer that such process could also have a role into the relationship between AD and other cancers. We thus quantified for each community of the -RBH AD/LC subnetwork the number of their connected pancancer metagenes (see Supposed Table 2 for results). Of note, the orientation of the TCGA pancancer metagenes has not been defined in [28], we thus cannot infer here if the activity of the pancancer metagenes is concordant with that of the LC metagenes or of the AD ones.

As reported in Supp Table 2, the majority of the identified communities, corresponding to the immune system-related signals, gender, chr χ and mitochondrial activity, matches metagenes obtained from other cancers, indicating a possible role of such processes in the co-morbidity of AD with other cancers. On the opposite five communities (19%), corresponding to LC subtypes, INF-Gamma and phosphate metabolism, are found to be specific to AD-LC inverse comorbidity.

## 3. Discussion

Matrix Factorization (MF) is a prominent solution for high-dimensional omics data analysis with a vast range of applications in computational biology.

We were here interested in investigating disease-disease relationships, representing an unprecedented opportunity to exploit mechanistic knowledge and repurpose treatments from one disease to the other. We thus proposed a computational framework for the application of MF to the study of disease-disease relationships. Considering the inverse comorbidity between Lung Cancer (LC) and Alzheimer’s Disease (AD) as a case study, different methodologies for the orientation of the metagenes were tested and the “Disease-pointing” one, orientating metagenes based on the case vs. control behavior of metasamples, proved to give better performance.

The framework here proposed and applied to the study of the inverse comorbidity between LC and AD, can be used to investigate direct/inverse comorbidity relations among other combinations of diseases. More complex patterns of direct and/or inverse comorbidities, involving more than two diseases, could also be studied. Moreover, we here chose sICA as MF algorithm and we employed transcriptomic data. However, the framework here proposed can also be implemented with other MF approaches (e.g. NMF, PCA) or different omics data types (e.g. methylome, proteome). More generally, also multi-omics factors, obtained with approaches such as Multi-Omics Factor Analysis (MOFA) or tensorial ICA (tICA), could also be considered as input of our analysis [58,59].

Finally, we performed a functional analysis of the genes involved in the subnetwork containing negative links between AD and LC factors. Our results confirmed previously identified molecular mechanisms underlying this inverse comorbidity, such as the involvement of the immune system and mitochondrial processes, plus new candidate factors have been identified. Overall, our results suggest that the MF RBH-based extended approach can be of biological and medical relevance once investigating the molecular bases of DD relationships.

## 4. Materials and Methods

### 4.1 Data collection

3 microarray datasets from NCBI Gene Expression Omnibus (GEO) (https://www.ncbi.nlm.nih.gov/geo/) were collected for Alzheimer’s disease: GSE4757; GSE48350 obtained from 4 brain regions: hippocampus, entorhinal cortex, superior frontal cortex, post-central gyrus and GSE5281 obtained from six brain regions: entorhinal cortex, hippocampus, medial temporal gyrus, posterior cingulate, superior frontal gyrus and primary visual cortex. The last two datasets were split based on the region of the brain in which the samples were collected, obtaining a total of 11 AD datasets composed of both case and control samples. Concerning lung cancer, 3 microarray datasets from NCBI GEO were collected: GSE19188, GSE19804 and GSE33532. The last one, involving 4 biopsies from the same sample, was split in 4 datasets. We thus obtained a total of 6 LC datasets composed of case and control samples. Additionally, the RNA-seq Lung dataset downloaded from The Cancer Genome Atlas (TCGA; https://tcga-data.nci.nih.gov/tcga/) was added to the analysis.

### 4.2 Biological characterization of the communities

We characterized the communities obtained in the -RBH AD/LC subnetwork using the following annotations: MSigDB signatures [33], Microenvironment Cell Populations-counter (MCP-counter) signatures [34], predefined TCGA lung cancer subtypes [35] and the metagenes computed in [27], here referred to as CIT. Concerning subtypes association we employed the metasamples obtained from the TCGA lung cancer data.

We tested the significance of the association with the predefined LC subtypes by performing a two-sided Wilcoxon test (cancer subtype vs. all other samples) and corrected for multiple testing using Bonferroni. For all the other biological annotations involving genes we employed the metagenes contained in each community. We associated to each community of the -RBH AD/LC subnetwork a “consensus metagene” corresponding to the average of all the metagenes contained in the community, paying attention to first concordantly orientate all the metagenes of the community based on the signs of their correlations (all the metagenes in the community were oriented based on the direction of LC). We then defined as top-contributing genes of a community those genes having a weight in the consensus metagene higher than 3 standard deviations in absolute value. The top-contributing genes were then divided into up and down based on their sign in the consensus metagene and tested for their intersection with the various collections of signatures. For cell types specific signatures we used a Fisher’s exact test with Bonferroni correction, for MsigDB we employed its default enrichment test [33].

After testing the association of each community with all the considered annotations (MSigDB signatures, MCP counter cell types signatures, the lung cancer subtypes available for TCGA data and the CIT metagenes), we associate to each community the annotation that is more consistently found across the different tests. In Supp Table 2 the annotations associated to each community together with their associated P-values are more extensively described.

Finally, to test the reproducibility of the identified consensus metagenes in other cancers, we used the metagenes computed with sICA on TCGA transcriptomics data from 32 different cancer types [28]. Then, for each community in the -RBH AD/LC subnetwork, we computed the number of cancers having at least one correlated metagene. The resulting values are reported in Supp Table 2.

## Supplementary Materials

Supplementary materials can be found at www.mdpi.com/xxx/s1.

## Author Contributions

conceptualization, L.C. and A.G.; methodology, L.C. and A.G.; formal analysis, L.C. and A.G.; biological curation, J.S, V.P, A.B. and A.V.; writing—original draft preparation, L.C. and A.B.; writing—review and editing, L.C., V.P., A.B.; supervision, L.C.; project administration, L.C.; funding acquisition, E.B. and A.Z.

## Funding

This work is supported by the “Departments of Excellence 2018–2022” Grant awarded by the Italian Ministry of Education, University and Research (MIUR) (L.232/2016). This work is also supported by a PhD Fellowship (BES-2016-077403) and funded by the Spanish Ministry of Economics and Competitiveness (BFU2015-71241-R).

## Acknowledgments

We thank the Bioinfo4Women program for partially funding V.P. and A.B. The results shown here are in part based upon data generated by the TCGA Research Network: https://www.cancer.gov/tcga.

## Conflicts of Interest

The authors declare no conflict of interest.

## Abbreviations

AD: Alzheimer’s disease
LC: Lung Cancer
RBH: Reciprocal Best Hit
DD: Disease-Disease

---

**Supp. Table 1.**
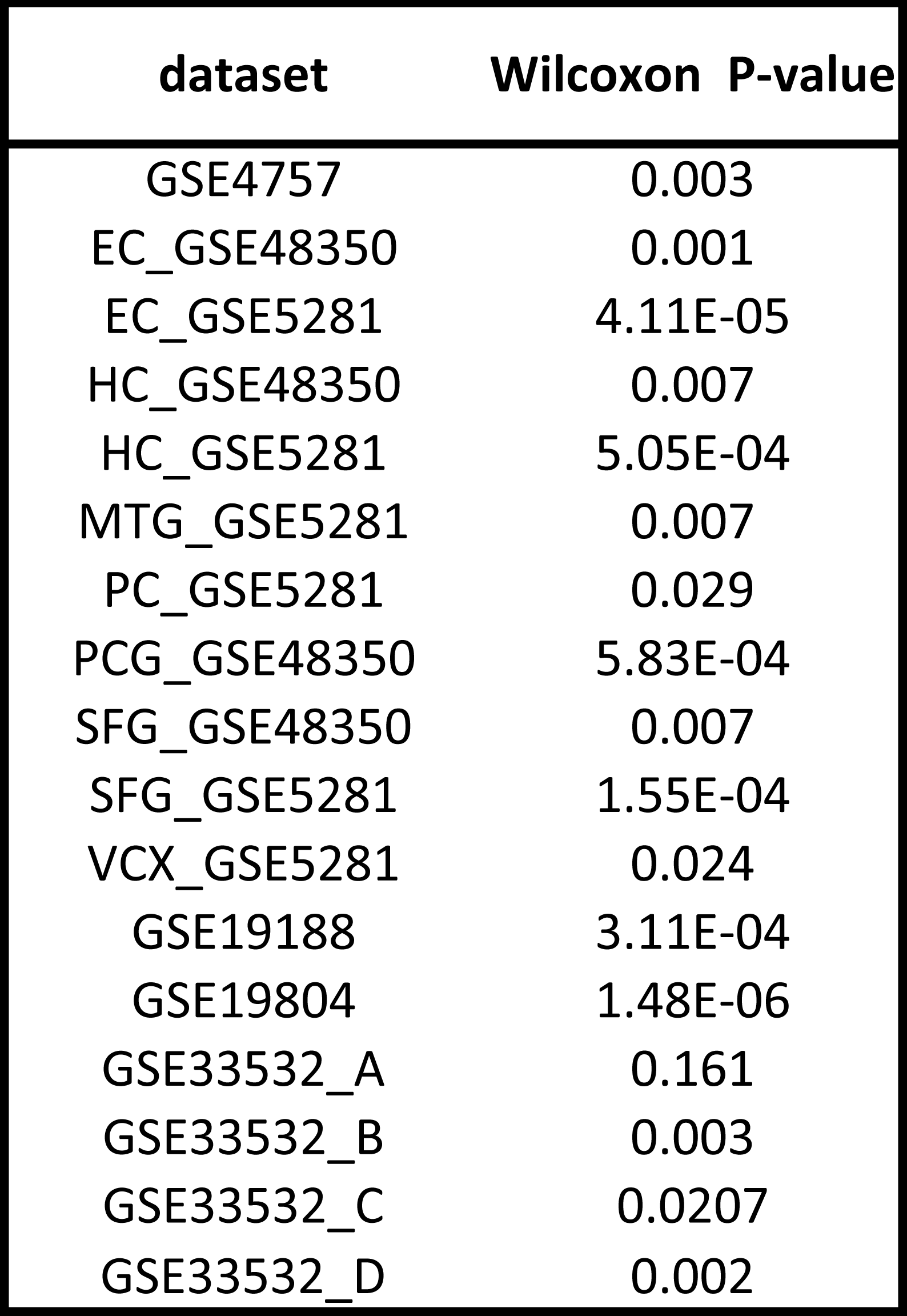
Comparison of correlation values between fold change of expression and metagenes oriented with long-tail vs disease-pointing approach.

| dataset | Wilcoxon P-value |
| --- | --- |
| GSE4757 | 0.003 |
| EC_GSE48350 | 0.001 |
| EC_GSE5281 | 4.11E-05 |
| HC_GSE48350 | 0.007 |
| HC_GSE5281 | 5.05E-04 |
| MTG_GSE5281 | 0.007 |
| PC_GSE5281 | 0.029 |
| PCG_GSE48350 | 5.83E-04 |
| SFG_GSE48350 | 0.007 |
| SFG_GSE5281 | 1.55E-04 |
| VCX_GSE5281 | 0.024 |
| GSE19188 | 3.11E-04 |
| GSE19804 | 1.48E-06 |
| GSE33532_A | 0.161 |
| GSE33532_B | 0.003 |
| GSE33532_C | 0.0207 |
| GSE33532_D | 0.002 |

**Supp. Table 2.**
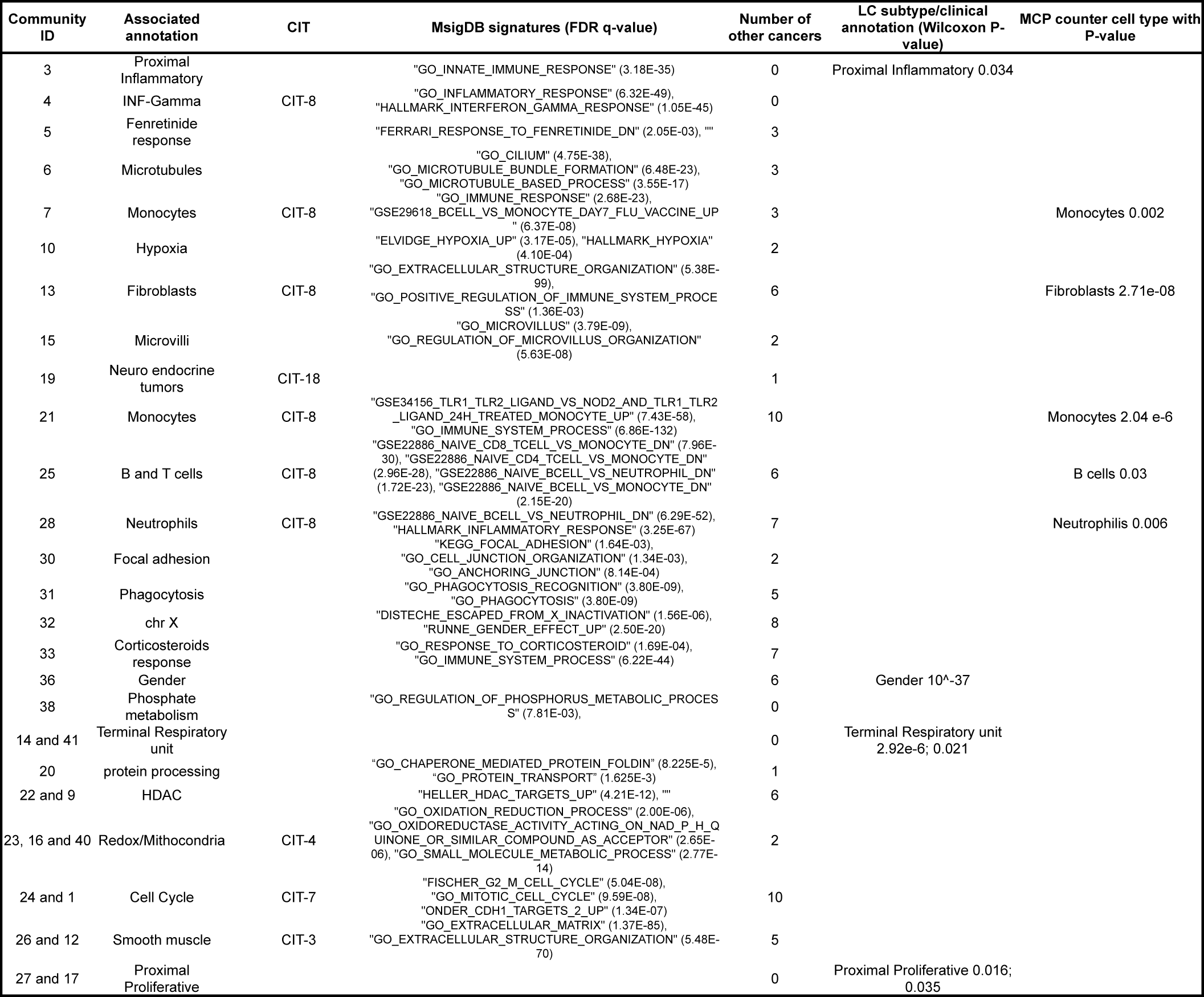
Annotations of the communities in the -RBH AD-LC subnetwork. For each community the table reports the ID, the associated annotation reported in the Figure, the correlated CIT metagene, the MsigDB enriched signatures with the associated FDR q-values, the number of TCGA cancer having a metagene correlated with them, the lung cancer subtype/clinical annotation with the Bonferroni corrected Wilcoxon P-value and the MCP counter based cell type with the Bonferroni corrected Fisher’s exact test P-value.

**Supp. Table 3.**
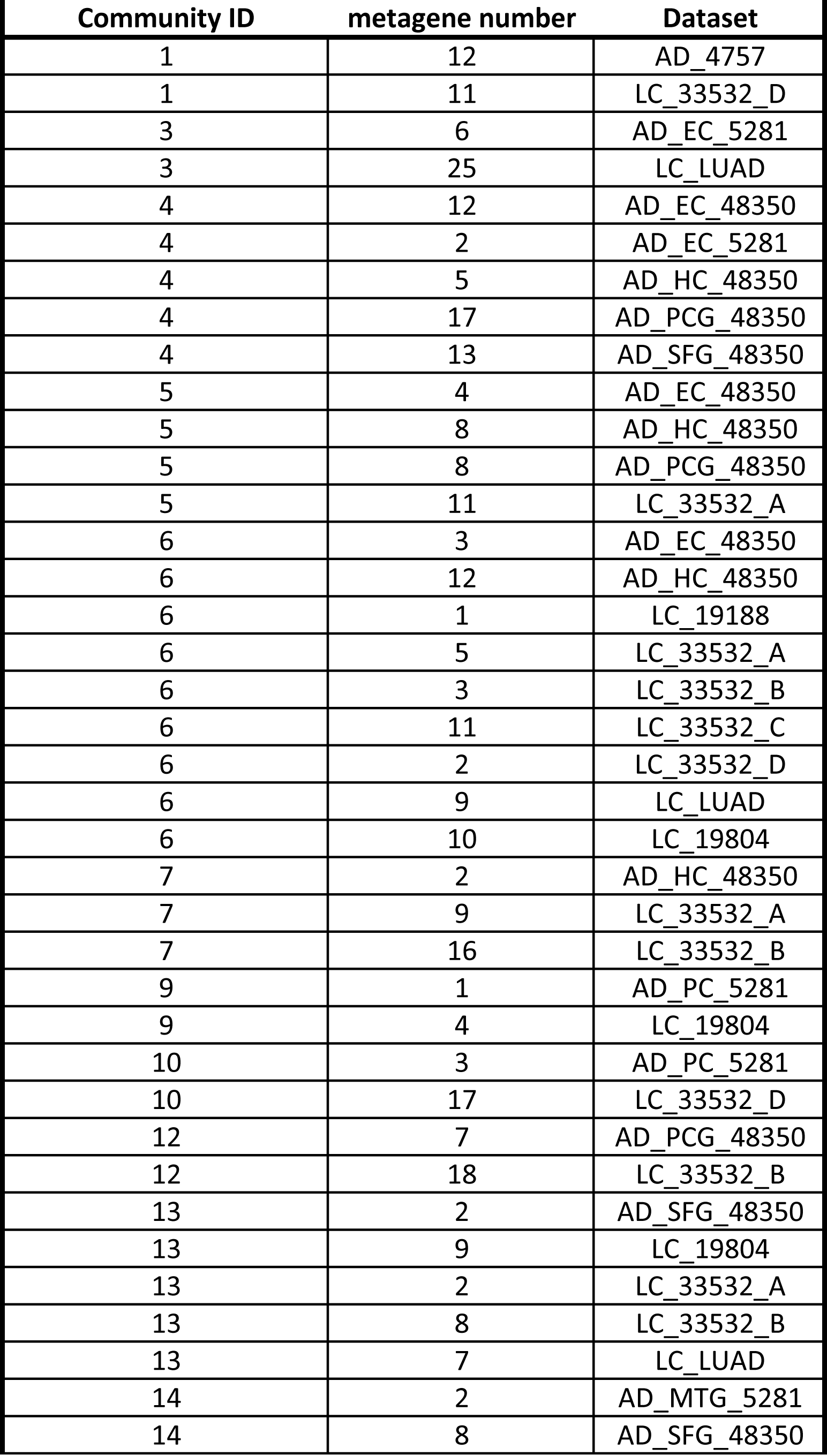
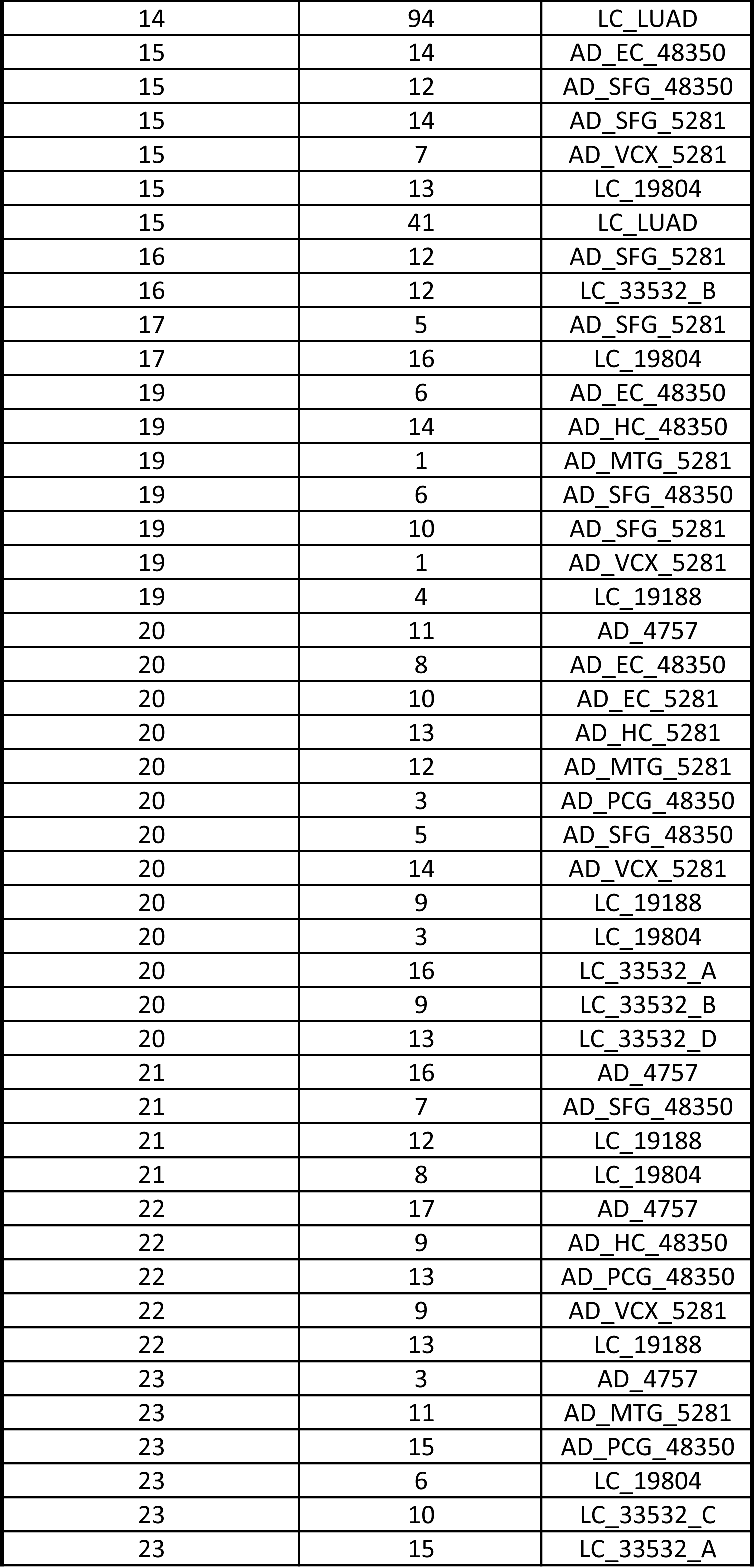
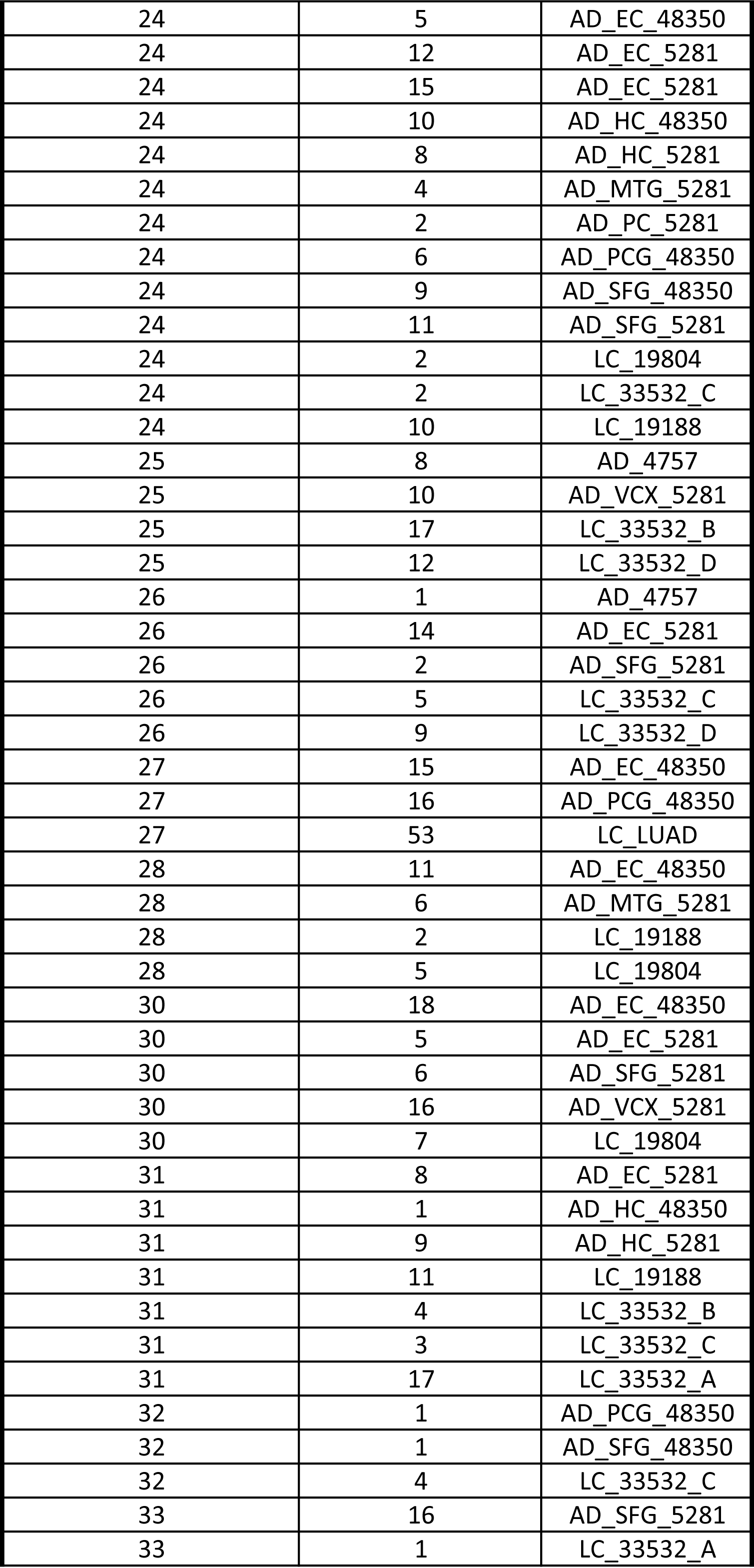
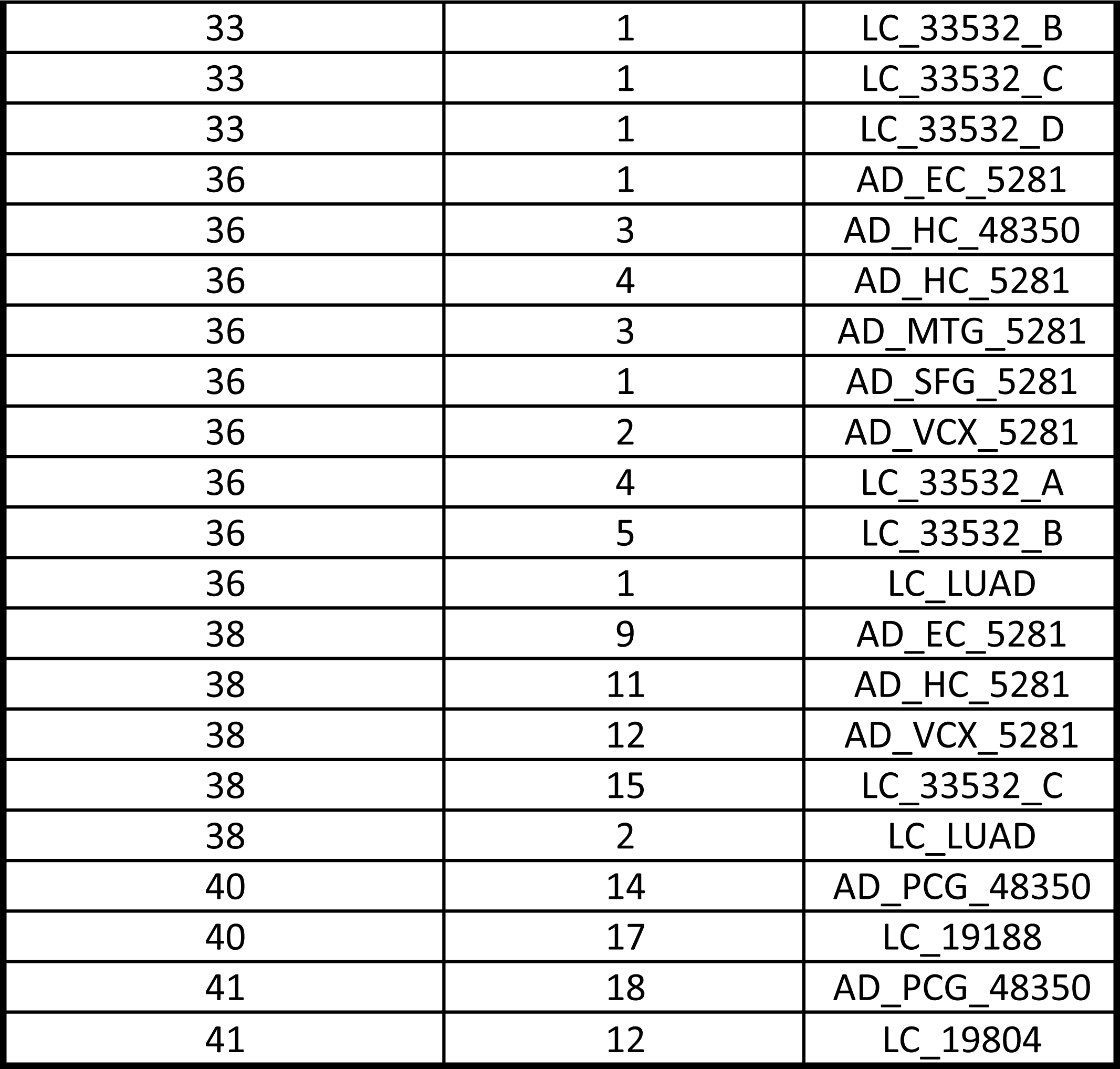
Annotation of the nodes present in the network of Figure 3. For each community ID reported in the figure, the components and associated datasets of its nodes are here reported corresponding to the

## References

1. Bell, G.; Hey, T.; Szalay, A. COMPUTER SCIENCE: Beyond the Data Deluge. Science 2009, 323, 1297–1298.

2. Stein-O’Brien, G.L.; Arora, R.; Culhane, A.C.; Favorov, A.V.; Garmire, L.X.; Greene, C.S.; Goff, L.A.; Li, Y.; Ngom, A.; Ochs, M.F.; et al. Enter the Matrix: Factorization Uncovers Knowledge from Omics. Trends Genet. 2018, 34, 790–805.

3. Devarajan, K. Nonnegative Matrix Factorization: An Analytical and Interpretive Tool in Computational Biology. PLoS Computational Biology 2008, 4, e1000029.

4. Meng, C.; Zeleznik, O.A.; Thallinger, G.G.; Kuster, B.; Gholami, A.M.; Culhane, A.C. Dimension reduction techniques for the integrative analysis of multi-omics data. Briefings in Bioinformatics 2016, 17, 628–641.

5. De Sousa E Melo, F.; Wang, X.; Jansen, M.; Fessler, E.; Trinh, A.; de Rooij, L.P.M.H.; de Jong, J.H.; de Boer, O.J.; van Leersum, R.; Bijlsma, M.F.; et al. Poor-prognosis colon cancer is defined by a molecularly distinct subtype and develops from serrated precursor lesions. Nature Medicine 2013, 19, 614–618.

6. Sadanandam, A.; Lyssiotis, C.A.; Homicsko, K.; Collisson, E.A.; Gibb, W.J.; Wullschleger, S.; Ostos, L.C.G.; Lannon, W.A.; Grotzinger, C.; Del Rio, M.; et al. A colorectal cancer classification system that associates cellular phenotype and responses to therapy. Nat. Med. 2013, 19, 619–625.

7. Kong, W.; Mou, X.; Hu, X. Exploring matrix factorization techniques for significant genes identification of Alzheimer’s disease microarray gene expression data. BMC Bioinformatics 2011, 12 Suppl 5, S7.

8. Brunet, J.-P.; Tamayo, P.; Golub, T.R.; Mesirov, J.P. Metagenes and molecular pattern discovery using matrix factorization. Proceedings of the National Academy of Sciences 2004, 101, 4164–4169.

9. Alexandrov, L.B.; Ju, Y.S.; Haase, K.; Van Loo, P.; Martincorena, I.; Nik-Zainal, S.; Totoki, Y.; Fujimoto, A.; Nakagawa, H.; Shibata, T.; et al. Mutational signatures associated with tobacco smoking in human cancer. Science 2016, 354, 618–622.

10. Australian Pancreatic Cancer Genome Initiative; ICGC Breast Cancer Consortium; ICGC MMML-Seq Consortium; ICGC PedBrain; Alexandrov, L.B.; Nik-Zainal, S.; Wedge, D.C.; Aparicio, S.A.J.R.; Behjati, S.; Biankin, A.V.; et al. Signatures of mutational processes in human cancer. Nature 2013, 500, 415–421.

11. Hackl, H.; Charoentong, P.; Finotello, F.; Trajanoski, Z. Computational genomics tools for dissecting tumour-immune cell interactions. Nat. Rev. Genet. 2016, 17, 441–458.

12. Cantini, L.; Kairov, U.; de Reyniès, A.; Barillot, E.; Radvanyi, F.; Zinovyev, A. Assessing reproducibility of matrix factorization methods in independent transcriptomes. Bioinformatics 2019.

13. Goh, K.-I.; Cusick, M.E.; Valle, D.; Childs, B.; Vidal, M.; Barabási, A.-L. The human disease network. Proc. Natl. Acad. Sci. U.S.A. 2007, 104, 8685–8690.

14. Halu, A.; De Domenico, M.; Arenas, A.; Sharma, A. The multiplex network of human diseases. NPJ Syst Biol Appl 2019, 5, 15.

15. Hidalgo, C.A.; Blumm, N.; Barabási, A.-L.; Christakis, N.A. A dynamic network approach for the study of human phenotypes. PLoS Comput. Biol. 2009, 5, e1000353.

16. Beck, M.K.; Jensen, A.B.; Nielsen, A.B.; Perner, A.; Moseley, P.L.; Brunak, S. Diagnosis trajectories of prior multi-morbidity predict sepsis mortality. Sci Rep 2016, 6, 36624.

17. Wang, K.; Gaitsch, H.; Poon, H.; Cox, N.J.; Rzhetsky, A. Classification of common human diseases derived from shared genetic and environmental determinants. Nat. Genet. 2017, 49, 1319–1325.

18. Eibl, G.; Cruz-Monserrate, Z.; Korc, M.; Petrov, M.S.; Goodarzi, M.O.; Fisher, W.E.; Habtezion, A.; Lugea, A.; Pandol, S.J.; Hart, P.A.; et al. Diabetes Mellitus and Obesity as Risk Factors for Pancreatic Cancer. J Acad Nutr Diet 2018, 118, 555–567.

19. Qu, Y.-L.; Liu, J.; Zhang, L.-X.; Wu, C.-M.; Chu, A.-J.; Wen, B.-L.; Ma, C.; Yan, X.-Y.; Zhang, X.; Wang, D.-M.; et al. Asthma and the risk of lung cancer: a meta-analysis. Oncotarget 2017, 8, 11614–11620.

20. Musicco, M.; Adorni, F.; Di Santo, S.; Prinelli, F.; Pettenati, C.; Caltagirone, C.; Palmer, K.; Russo, A. Inverse occurrence of cancer and Alzheimer disease: a population-based incidence study. Neurology 2013, 81, 322–328.

21. Freedman, D.M.; Wu, J.; Chen, H.; Kuncl, R.W.; Enewold, L.R.; Engels, E.A.; Freedman, N.D.; Pfeiffer, R.M. Associations between cancer and Alzheimer’s disease in a U.S. Medicare population. Cancer Med 2016, 5, 2965–2976.

22. Driver, J.A.; Beiser, A.; Au, R.; Kreger, B.E.; Splansky, G.L.; Kurth, T.; Kiel, D.P.; Lu, K.P.; Seshadri, S.; Wolf, P.A. Inverse association between cancer and Alzheimer’s disease: results from the Framingham Heart Study. BMJ 2012, 344, e1442.

23. Tavares, A.R.; de Melo, A.C.; Sternberg, C. Cancer linked to Alzheimer disease but not vascular dementia. Neurology 2010, 75, 1215–1216; author reply 1216.

24. Ganguli, M. A reduced risk of Alzheimer’s disease in those who survive cancer. BMJ 2012, 344, e1662.

25. Ibáñez, K.; Boullosa, C.; Tabarés-Seisdedos, R.; Baudot, A.; Valencia, A. Molecular evidence for the inverse comorbidity between central nervous system disorders and cancers detected by transcriptomic meta-analyses. PLoS Genet. 2014, 10, e1004173.

26. Sánchez-Valle, J.; Tejero, H.; Ibáñez, K.; Portero, J.L.; Krallinger, M.; Al-Shahrour, F.; Tabarés-Seisdedos, R.; Baudot, A.; Valencia, A. A molecular hypothesis to explain direct and inverse co-morbidities between Alzheimer’s Disease, Glioblastoma and Lung cancer. Sci Rep 2017, 7, 4474.

27. Biton, A.; Bernard-Pierrot, I.; Lou, Y.; Krucker, C.; Chapeaublanc, E.; Rubio-Pérez, C.; López-Bigas, N.; Kamoun, A.; Neuzillet, Y.; Gestraud, P.; et al. Independent component analysis uncovers the landscape of the bladder tumor transcriptome and reveals insights into luminal and basal subtypes. Cell Rep 2014, 9, 1235–1245.

28. Kairov, U.; Cantini, L.; Greco, A.; Molkenov, A.; Czerwinska, U.; Barillot, E.; Zinovyev, A. Determining the optimal number of independent components for reproducible transcriptomic data analysis. BMC Genomics 2017, 18, 712.

29. Engreitz, J.M.; Daigle, B.J.; Marshall, J.J.; Altman, R.B. Independent component analysis: Mining microarray data for fundamental human gene expression modules. Journal of Biomedical Informatics 2010, 43, 932–944.

30. Hyvärinen, A.; Oja, E. Independent component analysis: algorithms and applications. Neural Networks 2000, 13, 411–430.

31. van Dongen, S.; Abreu-Goodger, C. Using MCL to extract clusters from networks. Methods Mol. Biol. 2012, 804, 281–295.

32. Enright, A.J.; Van Dongen, S.; Ouzounis, C.A. An efficient algorithm for large-scale detection of protein families. Nucleic Acids Res. 2002, 30, 1575–1584.

33. Liberzon, A.; Subramanian, A.; Pinchback, R.; Thorvaldsdottir, H.; Tamayo, P.; Mesirov, J.P. Molecular signatures database (MSigDB) 3.0. Bioinformatics 2011, 27, 1739–1740.

34. Becht, E.; Giraldo, N.A.; Lacroix, L.; Buttard, B.; Elarouci, N.; Petitprez, F.; Selves, J.; Laurent-Puig, P.; Sautès-Fridman, C.; Fridman, W.H.; et al. Estimating the population abundance of tissue-infiltrating immune and stromal cell populations using gene expression. Genome Biology 2016, 17.

35. Cancer Genome Atlas Research Network Comprehensive molecular profiling of lung adenocarcinoma. Nature 2014, 511, 543–550.

36. Beeri, M.S.; Schmeidler, J.; Lesser, G.T.; Maroukian, M.; West, R.; Leung, S.; Wysocki, M.; Perl, D.P.; Purohit, D.P.; Haroutunian, V. Corticosteroids, but not NSAIDs, are associated with less Alzheimer neuropathology. Neurobiol. Aging 2012, 33, 1258–1264.

37. Scott, S.C.; Pennell, N.A. Early Use of Systemic Corticosteroids in Patients with Advanced NSCLC Treated with Nivolumab. J Thorac Oncol 2018, 13, 1771–1775.

38. Ohlmann, C.-H.; Jung, C.; Jaques, G. Is growth inhibition and induction of apoptosis in lung cancer cell lines by fenretinide [N-(4-hydroxyphenyl)retinamide] sufficient for cancer therapy? Int. J. Cancer 2002, 100, 520–526.

39. Goodman, A.B. Retinoid receptors, transporters, and metabolizers as therapeutic targets in late onset Alzheimer disease. J. Cell. Physiol. 2006, 209, 598–603.

40. Lin, C.-H.; Lee, S.-Y.; Zhang, C.-C.; Du, Y.-F.; Hung, H.-C.; Wu, H.-T.; Ou, H.-Y. Fenretinide inhibits macrophage inflammatory mediators and controls hypertension in spontaneously hypertensive rats via the peroxisome proliferator-activated receptor gamma pathway. Drug Des Devel Ther 2016, 10, 3591–3597.

41. Peers, C.; Pearson, H.A.; Boyle, J.P. Hypoxia and Alzheimer’s disease. Essays Biochem. 2007, 43, 153–164.

42. Nalivaeva, N.N.; Turner, A.J.; Zhuravin, I.A. Role of Prenatal Hypoxia in Brain Development, Cognitive Functions, and Neurodegeneration. Frontiers in Neuroscience 2018, 12.

43. Salem, A.; Asselin, M.-C.; Reymen, B.; Jackson, A.; Lambin, P.; West, C.M.L.; O’Connor, J.P.B.; Faivre-Finn, C. Targeting Hypoxia to Improve Non-Small Cell Lung Cancer Outcome. J. Natl. Cancer Inst. 2018, 110.

44. Mazure, C.M.; Swendsen, J. Sex differences in Alzheimer’s disease and other dementias. Lancet Neurol 2016, 15, 451–452.

45. Zang, E.A.; Wynder, E.L. Differences in Lung Cancer Risk Between Men and Women: Examination of the Evidence. JNCI Journal of the National Cancer Institute 1996, 88, 183–192.

46. Patra, S.; Panigrahi, D.P.; Praharaj, P.P.; Bhol, C.S.; Mahapatra, K.K.; Mishra, S.R.; Behera, B.P.; Jena, M.; Bhutia, S.K. Dysregulation of histone deacetylases in carcinogenesis and tumor progression: a possible link to apoptosis and autophagy. Cell. Mol. Life Sci. 2019.

47. Janczura, K.J.; Volmar, C.-H.; Sartor, G.C.; Rao, S.J.; Ricciardi, N.R.; Lambert, G.; Brothers, S.P.; Wahlestedt, C. Inhibition of HDAC3 reverses Alzheimer’s disease-related pathologies in vitro and in the 3xTg-AD mouse model. Proceedings of the National Academy of Sciences 2018, 115, E11148–E11157.

48. Zhang, H.; Shao, H.; Golubovskaya, V.M.; Chen, H.; Cance, W.; Adjei, A.A.; Dy, G.K. Efficacy of focal adhesion kinase inhibition in non-small cell lung cancer with oncogenically activated MAPK pathways. Br. J. Cancer 2016, 115, 203–211.

49. Lachén-Montes, M.; González-Morales, A.; de Morentin, X.M.; Pérez-Valderrama, E.; Ausín, K.; Zelaya, M.V.; Serna, A.; Aso, E.; Ferrer, I.; Fernández-Irigoyen, J.; et al. An early dysregulation of FAK and MEK/ERK signaling pathways precedes the β-amyloid deposition in the olfactory bulb of APP/PS1 mouse model of Alzheimer’s disease. J Proteomics 2016, 148, 149–158.

50. Fuchsberger, T.; Martínez-Bellver, S.; Giraldo, E.; Teruel-Martí, V.; Lloret, A.; Viña, J. Aβ Induces Excitotoxicity Mediated by APC/C-Cdh1 Depletion That Can Be Prevented by Glutaminase Inhibition Promoting Neuronal Survival. Sci Rep 2016, 6, 31158.

51. Yu, Q.; Guo, Q.; Chen, L.; Liu, S. Clinicopathological significance and potential drug targeting of CDH1 in lung cancer: a meta-analysis and literature review. Drug Des Devel Ther 2015, 9, 2171–2178.

52. Ashraf, G.M.; Greig, N.H.; Khan, T.A.; Hassan, I.; Tabrez, S.; Shakil, S.; Sheikh, I.A.; Zaidi, S.K.; Akram, M.; Jabir, N.R.; et al. Protein misfolding and aggregation in Alzheimer’s disease and type 2 diabetes mellitus. CNS Neurol Disord Drug Targets 2014, 13, 1280–1293.

53. Selkoe, D.J. Cell biology of protein misfolding: the examples of Alzheimer’s and Parkinson’s diseases. Nat. Cell Biol. 2004, 6, 1054–1061.

54. Vallin, J.; Grantham, J. The role of the molecular chaperone CCT in protein folding and mediation of cytoskeleton-associated processes: implications for cancer cell biology. Cell Stress P. Chaperones 2019, 24, 17–27.

55. Slooter, A.J.; Bronzova, J.; Witteman, J.C.; Van Broeckhoven, C.; Hofman, A.; van Duijn, C.M. Estrogen use and early onset Alzheimer’s disease: a population-based study. J. Neurol. Neurosurg. Psychiatry 1999, 67, 779–781.

56. Vegeto, E.; Benedusi, V.; Maggi, A. Estrogen anti-inflammatory activity in brain: a therapeutic opportunity for menopause and neurodegenerative diseases. Front Neuroendocrinol 2008, 29, 507–519.

57. Rodriguez-Lara, V.; Hernandez-Martinez, J.-M.; Arrieta, O. Influence of estrogen in non-small cell lung cancer and its clinical implications. J Thorac Dis 2018, 10, 482–497.

58. Argelaguet, R.; Velten, B.; Arnol, D.; Dietrich, S.; Zenz, T.; Marioni, J.C.; Buettner, F.; Huber, W.; Stegle, O. Multi-Omics Factor Analysis—a framework for unsupervised integration of multiomics data sets. Molecular Systems Biology 2018, 14, e8124.

59. Teschendorff, A.E.; Jing, H.; Paul, D.S.; Virta, J.; Nordhausen, K. Tensorial blind source separation for improved analysis of multi-omic data. Genome Biology 2018, 19.

